# Emotion lies in the eye of the listener: emotional arousal to novel sounds is reflected in the sympathetic contribution to the pupil dilation response and the P3

**DOI:** 10.1101/250084

**Authors:** Andreas Widmann, Erich Schröger, Nicole Wetzel

**Author notes:** Corresponding author: Andreas Widmann, Institute of Psychology, University of Leipzig, Neumarkt 9-19, D-04109 Leipzig Germany.

## Abstract

Novel sounds in the auditory oddball paradigm elicit a biphasic dilation of the pupil (PDR) and P3a as well as novelty P3 event-related potentials (ERPs). The biphasic PDR has been hypothesized to reflect the relaxation of the iris sphincter muscle due to parasympathetic inhibition and the constriction of the iris dilator muscle due to sympathetic activation. We measured the PDR and the P3 to neutral and to emotionally arousing negative novels in dark and moderate lighting conditions. By means of principal component analysis (PCA) of the PDR data we extracted two components: the early one was absent in darkness and, thus, presumably reflects parasympathetic inhibition, whereas the late component occurred in darkness and light and presumably reflects sympathetic activation. Importantly, only this sympathetic late component was enhanced for emotionally arousing (as compared to neutral) sounds supporting the hypothesis that emotional arousal specifically activates the sympathetic nervous system. In the ERPs we observed P3a and novelty P3 in response to novel sounds. Both components were enhanced for emotionally arousing (as compared to neutral) novels. Our results demonstrate that sympathetic and parasympathetic contributions to the PDR can be separated and link emotional arousal to sympathetic nervous system activation.

**Highlights:**

- PDR and ERP effects of novel emotional and neutral oddball sounds were studied.
- Parasympathetic and sympathetic contributions to the PDR were dissociated by PCA.
- The parasympathetic PDR component was absent in darkness.
- Emotional arousal enhanced the sympathetic contribution to the PDR and the P3 ERP.
- Effects of emotional arousal are mediated by the sympathetic pathway.

## Introduction

The pupil diameter is controlled by both autonomic nervous systems via two muscles. The ring-shaped iris sphincter muscle is innervated by the parasympathetic nervous system while the radial iris dilator muscle is innervated by the sympathetic nervous system (Einhäuser, 2017; McDougal & Gamlin, 2008). Both nervous systems closely couple the pupil to the locus coeruleus-norepinephrine (LC-NE) system (via an inhibitory projection to the Edinger-Westphal nucleus in case of the parasympathetic system). The LC-NE system is a neuromodulatory system that is central for various functions, in particular attention, emotion, motivation, decision making, and memory (Aston-Jones & Cohen, 2005; Murphy, O'Connell, O'Sullivan, Robertson, & Balsters, 2014; Murphy, Robertson, Balsters, & O'Connell, 2011; Preuschoff, ’t Hart, & Einh#x00E4;user, 2011; for review see, Laeng, Sirois, & Gredebäck, 2012; Einhäuser, 2017). The LC-NE system shows a phasic activation in response to task-relevant, salient, or arousing stimuli (for review see e.g.,Sara & Bouret, 2012). Rare or novel events presented within a sequence of frequently repeated standard events (the oddball paradigm) do elicit a pupil dilation response (PDR; Friedman, Hakerem, Sutton, & Fleiss, 1973; Hochmann & Papeo, 2014; Liao, Kidani, Yoneya, Kashino, & Furukawa, 2016; Liao, Yoneya, Kidani, Kashino, & Furukawa, 2016; Murphy et al., 2014; Murphy et al., 2011; Qiyuan, Richer, Wagoner, & Beatty, 1985; Steinhauer & Hakerem, 1992; Wetzel, Buttelmann, Schieler, & Widmann, 2016).

Independently from sounds’ novelty, the PDR has also been shown to be highly sensitive to emotional arousal. For example, Partala and Surakka (2003) reported enhanced PDRs in response to sounds with positive or negative valence compared to neutral sounds. Also in the visual modality, Bradley, Miccoli, Escrig, and Lang (2008) observed larger pupil dilation responses for pleasant and unpleasant emotionally arousing pictures compared to emotionally neutral pictures. In their study Bradley and colleagues observed a co-variation of the PDR and skin conductance changes but no co-variation of PDR and cardiac deceleration. Based on this indirect evidence Bradley and colleagues (2008) suggested that the emotion-related PDR effects are supposed to be specific for sympathetic nervous system activity. However, this has not yet been directly shown on the basis of the PDR itself. Both nervous systems antagonistically contribute to the observed pupil dilation: relaxation of the iris sphincter muscle by parasympathetic inhibition and constriction of the iris dilator muscle by sympathetic activation. Attempts to non-invasively dissociate their contributions have not yet been successful (Einhäuser, 2017). The possibility to separate the differential contributions of the parasympathetic and sympathetic nervous systems to the PDR is therefore of high relevance.

Frequently, the observed pupil dilation in response to rare events is biphasic and it has been suggested that the two peaks reflect the overlapping contributions of the sphincter muscle relaxation by parasympathetic inhibition and of the dilator muscle constriction by sympathetic activation (Steinhauer & Hakerem, 1992). Steinhauer and Hakerem (1992) hypothesized that overlapping sympathetic and parasympathetic contributions should be separable by means of factor analysis. A biphasic response to several environmental distractor sounds was also observed in infants and adults (Wetzel et al., 2016). Two main components were extracted using a principal component analysis (PCA), however, the attribution of components to changes in parasympathetic and sympathetic activity remained unclear. A PCA was also applied to pupillary data by Geuter, Gamer, Onat, and Büchel (2014). The components were, however, not related to pupil physiology but used to predict pain ratings by a regression based approach. Together with their original hypothesis Steinhauer and Hakerem (1992) already suggested a possible validation of this attribution by variation of lighting conditions: the parasympathetic tone, but not sympathetic tone, is minimal in darkness (Loewenfeld, 1958; and the sphincter muscle maximally relaxed). Thus, in darkness, a component related to parasympathetic inhibition should be reduced in comparison to bright lighting conditions, while a component related to sympathetic activation is not supposed to be modulated by lighting conditions. The contributions of the parasympathetic and the sympathetic pathway to pupil dilation during sustained processing have already been successfully separated by the variation of ambient lighting as well as by pharmacological manipulations (Steinhauer, Siegle, Condray, & Pless, 2004). However, for the phasic PDR observed in response to rare (sound) events the separation of the contributions of parasympathetic inhibition and sympathetic activation by means of factor analysis or PCA including a validation by lighting conditions has-to our knowledge-not yet been empirically tested.

Also the P3-family of event-related potential (ERP) responses associated with processes of attention and memory (for review see, e.g., Alho, Escera, & Schröger, 2003; Polich, 2007) has been suggested to be related to LC-NE system activity. More specifically, a current theory postulates that P3 and the autonomic component of the orienting response do reflect the co120 activation of the locus coeruleus-norepinephrine (LC-NE) system and the peripheral sympathetic nervous system (SNS; Nieuwenhuis, De Geus, & Aston-Jones, 2011). The P3a subcomponent is elicited by unexpected stimuli that violate a regularity previously built up on the basis of the sensory input. Similar to the PDR, the P3a can be elicited by salient, arousing, rare, novel, or task-relevant events presented amongst regular standard stimuli in an oddball paradigm. In response to novel stimuli frequently a double peaked P3 component was observed and reported in the literature as P3a and novelty P3 (Barry, Steiner, & De Blasio, 2016; Friedman, Cycowicz, & Gaeta, 2001) or early and late P3a (Escera, Alho, Winkler, & Näätänen, 1998; Yago, Escera, Alho, Giard, & Serra-Grabulosa, 2003). P3a (early P3a) and novelty P3 (late P3a) are supposed to indicate orienting of attention and enhanced evaluation of significant or novel stimuli (Alho et al., 2003; Polich, 2007).

The P3a component has been shown to be enhanced or modulated in response to emotional compared to neutral auditory (Pakarinen et al., 2014; Thierry & Roberts, 2007; but also see, Czigler, Cox, Gyimesi, & Horvath, 2007, not finding an effect) and visual stimuli (e.g., Keil et al., 2002) indicating stronger orienting of attention. Also the P3b and the late positive/slow wave components have been reported to be sensitive to emotionally arousing stimuli (Cuthbert, Schupp, Bradley, Birbaumer, & Lang, 2000; Delplanque, Silvert, Hot, Rigoulot, & Sequeira, 2006; Delplanque, Silvert, Hot, & Sequeira, 2005; Foti, Hajcak, & Dien, 2009; Keil et al., 2002; for review see e.g., Bradley, Keil, & Lang, 2012). If emotional arousal modulates sympathetic nervous system activity a co-modulation of the sympathetic component of PDR and P3 would be expected and give some support to the suggestion that the P3-family of ERP components might reflect the co-activation of the LC-NE system and the sympathetic nervous system (even if PDR and P3 amplitude do not necessarily correlate at the single trial level; Kamp & Donchin, 2015).

Here we aimed at examining these issues in detail by the co-registration of the PDR and P3 in an auditory oddball paradigm including emotionally arousing negative and neutral novel sounds in dark and moderate lighting conditions. We hypothesized that we can replicate the decomposition of the biphasic PDR to novel sounds into two components by means of PCA (as already demonstrated by Wetzel et al., 2016), additionally confirming the parasympathetic vs. sympathetic origin of the two components by variation of lighting conditions. Further, we hypothesized that only the sympathetic contribution would be modulated by emotional arousal. Finally, following the LC-NE SNS co-activation hypothesis we expected an enhanced P3 in response to emotionally arousing novel sounds.

## Materials and Methods

### Participants

A total of 22 young adults participated in the experiment, 2 of them were excluded from data analysis due to excessive blinking resulting in less than 70% of trials remaining in any condition after EEG and pupil data artifact rejection. The exclusion criterion was defined a posteriori. A total of 13 of the participants included in data analysis were female, 7 male, 16 were right-handed, and 4 left-handed. Their mean age was 23 years and 10 months (range 18;0–38;8 years;months). Participation was rewarded by money or course credit points. Participants gave written informed consent and confirmed normal hearing abilities, normal or corrected-to-normal vision, and to be not under the influence of pharmacological substances that affect the central nervous system. The project was approved by the local ethics committee of the University.

### Stimuli

Novel sounds were 32 high arousing negative sounds and 32 moderately arousing neutral sounds. The standard sound was a sine wave sound with a fundamental frequency of 500 Hz including the second and third harmonic attenuated by –3 and –6 dB respectively. Sounds had a duration of 500 ms including faded ends of 5 ms. Sounds were present at a loudness level of 56.1 dB SPL measured with HMS III dummy head (HEAD Acoustics, Herzogenrath, Germany). Loudness was equalized with root mean square normalization.

The novel sounds were used and described in detail in a previous study by Max, Widmann, Kotz, Schröger, and Wetzel (2015). There, 32 high arousing emotionally negative and 32 moderately arousing emotionally neutral sounds were selected in a pilot study from a set of 210 sounds collected from the International Affective Digitized Sounds (IADS; Bradley & Lang, 2007) from a study by Hasting, Wassiliwizky, and Kotz (2010), and from other data bases as described by Max et al. (2015). Sounds were rated for valence (unpleasant – neutral – pleasant) and arousal (calm – arousing) on 9-point scales with the Self-Assessment Manikins (Bradley & Lang, 1994). Valence and arousal ratings were significantly different for emotionally negative vs. neutral sounds (Max et al., 2015).

The visual target consisted of two continuously presented, horizontally or vertically aligned grey single pixel dots (±0.12° to the left and right or above and below the center of the screen respectively). In the dark condition, visual dots were presented on black background (0 cd/m^2^ as measured by Cambridge Research Systems OptiCal and confirmed by Minolta LS-100 luminance meters) and had a luminance of 16.16 cd/m^2^. In t he light condition, visual dots were presented on dark grey background (0.75 cd/m^2^) and had a luminance of 18.97 cd/m^2^.

### Apparatus

Participants were seated in a dark, electrically shielded, double-walled sound booth (Industrial Acoustics Company). All possible light sources (display and EEG amplifier LEDs, etc.) except the CRT display and the eye-tracker IR LED panel were removed or covered. Participants’ heads were stabilized with a chin and forehead rest. Visual stimuli were presented on a 19” CRT display (resolution 1024 × 768, refresh rate 100 Hz, distance 60 cm; G90fB; ViewSonic). Display brightness was adjusted to the minimal possible value. The task of the participants was to track and count the number orientation changes of the visual target (per block). Sounds were presented with loudspeakers (Bose Companion 2 Series II, Bose Corporation, Framingham, MA) on the left and right of the display.

### Procedure

The dark and the light conditions each consisted of four blocks separated by short breaks without changes in ambient luminance. Between conditions participants had a longer resting period of several minutes at comfortable ambient lighting. Each condition started with a one-minute dark adaptation period without visual or auditory stimulation. Each block started with a nine-point eye-tracker calibration and validation procedure followed by an additional 15-second adaptation period without stimulation. The block duration including eye-tracker calibration and adaptation was approximately 6 min. In each block 144 sounds – 120 standard sounds (83.3%), 12 neutral novels (8.3%), and 12 emotional novels (8.3%) – were presented in pseudo-randomized order. At least two standard sounds were presented between two novels. Sixteen novels – randomly selected for each participant, condition, and category – were presented once, the other 16 novels twice per condition but never within the same block.

The SOA between sounds was 1.8, 2, 2.2, or 2.4 s with equal probability. The orientation changes of the visual target were Poisson distributed at a rate of 5 min^−1^. The delay between two target orientation changes was at least 0.6 s. The latency of target orientation changes was adjusted not to occur within ±0.2 s before or after the onset of a sound to avoid illusory audio217 visual events. Within a ±1.8 s period before and after each novel sound the probability of target orientation changes was reduced so that not more than one novel per category and block had to be discarded (resulting in slightly reduced overall rate). On average 20.5 target orientation changes occurred per block (range 10 to 34). Participants were asked to report the counted number of orientation changes after each block and to ignore the auditory stimulation. The experiment lasted about 2 hours including breaks, application of electrodes, and the adjustment of the eye-tracker.

### Data Recording

The pupil diameter of both eyes was recorded with an infrared EyeLink 1000 (SR Research, Ottawa, Ontario, Canada) remote eye tracker at a sampling rate of 500 Hz. The electroencephalogram (EEG) was recorded with a BrainAmp MR amplifier by Brain Products GmbH (Gilching, Germany) at a sampling rate of 500 Hz. 32 Ag/Ag-Cl-electrodes were placed at the following positions of the extended 10-20 system: Fp1, Fp2, F7, F3, Fz, F4, F8, FC5, FC1, FC2, FC6, T7, C3, Cz, C4, T8, CP5, CP1, CP2, CP6, P7, P3, Pz, P4, P8, O1, O2, and at the left (M1) and right (M2) mastoids. Three electrodes recording horizontal and vertical electrooculogram (EOG) components were positioned to the left and right of the outer canthi of the eyes and below the left eye. The reference electrode was placed at the tip of the nose.

### Pupil data preprocessing

Eye tracker pupil diameter digital counts were calibrated using the method suggested by Marchak and Steinhauer (2011) and converted to mm. Blinks were marked by the blink events provided by the eye-tracker (including the enclosing saccade event markers). Additionally, partial blinks (not reported by the eye-tracker) were detected from the smoothed velocity times series as pupil diameter changes exceeding 20 mm/s including a 50 ms pre-blink and a 100 ms post-blink interval (Merritt, Keegan, & Mercer, 1994). Blinks (and intervals with signal loss) longer than 1 s were discarded from the continuous data and from further analysis. Data were segmented into epochs of 2 s duration including a .2 s pre-stimulus baseline and baseline corrected by subtracting the mean amplitude of the baseline period from each epoch. The first two (standard) sounds per block and each two (standard) sound immediately following a deviant sound were removed from further analysis to avoid the contamination of responses to standards by deviant related effects (see e.g., Wetzel, 2015). All trials including an orientation change of the visual target from 1.8 s before to 1.8 s after sound onset were excluded from further analysis. All epochs including blinks accounting for more than 50% of epoch duration at any eye were excluded from further analysis. Only corresponding identical trials were included in both pupil and EEG data analysis. That is, trials excluded from any, pupil or EEG data, were excluded from both types of analyses. Individual average PDRs were computed per participant and condition from the mean of both eyes. Data during blinks were excluded from averaging. Grand-average waveforms were computed from the individual average PDRs for each lighting condition and sound type.

### Pupil data PCA analysis

We performed a temporal principal component analysis to separate the main factors underlying the biphasic response with the ERP PCA Toolkit MATLAB toolbox by Dien (2010). The PCA was computed on the individual average PDR data including all novel and lighting conditions. PCA was computed using Promax rotation (κ = 3) with a covariance relationship matrix and Kaiser weighting. Based on Horn’s parallel test two components were retained (explaining more than 95% of the variance). Note that components are ordered by explained variance and not chronological by peak latency (that is, component 2 peaks earlier than component 1).

### EEG data preprocessing

EEG data analysis was performed with MATLAB software and the EEGLAB toolbox (Delorme & Makeig, 2004). Data were filtered offline with a 0.1 Hz high269 pass filter (Kaiser windowed sinc FIR filter, order=8024, beta=5, transition band width=0.2 Hz; Widmann & Schröger, 2012; Widmann, Schröger, & Maess, 2015) and a 40 Hz low-pass filter (Kaiser windowed sinc FIR filter, order = 162, beta = 5, transition band width = 10 Hz.

The data were segmented into epochs of .8 s duration including a .2 s pre-stimulus baseline. Noisy channels with a robust z-score of the robust standard deviation larger than 3 were removed from the data (a single channel in 4 of the participants; Bigdely-Shamlo, Mullen, Kothe, Su, & Robbins, 2015). Data were corrected for artifacts using independent component analysis (ICA). To improve the decomposition the ICA was computed on the raw data (excluding bad channels) filtered by a 1 Hz high-pass filter (Kaiser windowed sinc FIR filter, order = 804, beta = 5, transition band width = 2 Hz) and a 40 Hz low-pass filter (see above) and segmented into epochs of 1.5 s duration (−0.5 to 1 s relative to sound onset; but not baseline corrected; Groppe, Makeig, & Kutas, 2009). The obtained demixing matrix was then applied to the 0.1–40 Hz filtered data. Winkler, Debener, Müller, and Tangermann (2015) have suggested and validated that high-pass filters do improve ICA decompositions (reliability, independence, dipolarity) and the demixing matrix can be applied to a linear transformed dataset. Artifact ICs were detected using the SASICA EEGLAB plugin including the low autocorrelation, focal channel or trial activity, correlation with vertical or horizontal EOG, and ADJUST criteria (Chaumon, Bishop, & Busch, 2015; Mognon, Jovicich, Bruzzone, & Buiatti, 2011) followed by additional manual IC classification. Artifact IC activity was subtracted from the data. On average 9.3 ICs were removed from the data per participant (median = 9.5; min = 6; max = 13). Subsequently, bad channels were interpolated using spherical spline interpolation. Data were baseline corrected by subtracting the mean amplitude of the baseline period from each epoch. Trials with peak-to-peak voltage differences exceeding 200 µV, the first two trials per block, two standard trials following a novel trial and trials excluded from pupil data were excluded from the analysis. Individual average ERPs were computed per participant and sound type and averaged over lighting conditions. We had no a priori hypothesis on the effects of lighting conditions on the ERPs and an analysis of lighting conditions would have required doubling the trial numbers and recording the experiment in two separate sessions. Grand-average waveforms were computed from the individual average ERPs for each sound type.

### ERP data PCA analysis

Analogous to the pupil data a temporal PCA was computed on the individual average ERP data including all channels and novel conditions with the same parameters as for the PDR analysis. Eight components were retained. The statistical analysis was restricted to components 1 and 3 reflecting P3a (early P3a) and novelty P3 (late P3a).

### Trial numbers

On average 208.6 standard (Mdn = 211.5; SD = 13.1; min = 174; max = 231), 42.1 emotional (Mdn = 43; SD = 2.6; min = 34; max = 44), and 42.8 neutral novel trials (Mdn = 44; SD = 2.5; min = 35; max = 45) in the dark condition and 206.2 standard (Mdn = 204; SD = 14.9; min = 177; max = 231), 42.9 emotional (Mdn = 44; SD = 1.7; min = 38; max = 44), and 43.2 neutral novel trials (Mdn = 44; SD = 1.9; min = 38; max = 45) in the light condition were included into the analysis per participant. The ratio of presented vs. included trials was not significantly different between conditions (*F*(5,95) = 0.773, *p* = .414, *ε* = .247, *η*_G_^2^ = .026). In total 340 trials (M = 17; Mdn = 13; SD = 15.8; min = 0; max = 62) were excluded due to blink periods longer than 1 s or accounting for more than half of the trial duration in the pupil data. In total 16 trials (M = .8; Mdn = 0; SD = 1.7; min = 0; max = 6) were excluded due to peak-to-peak voltage differences exceeding 200 µV in the EEG. In total 2 trials were excluded due to artifacts in pupil *and* EEG data. Blinks accounted for 4.1 % of measurement time (Mdn = 3 % SD = 3.3 %; min = 0.4 %; max = 10.7 %) and were not included into averaging. The proportion of blink periods was not significantly different between conditions (*F*(5,95) = 1.391, *p* = .26, *ε* = .288, *η*_G_^2^ = .016)

### Statistical analysis

PDR PCA component scores were tested with a repeated measures ANOVA including the within-subject factors *emotion* (emotionally negative vs. neutral) × *component* (2 vs. 1) × *lighting* (dark vs. light). ERP PCA component scores were tested with a repeated-measures ANOVA including the within-subject factors *emotion* × *component* (1 vs. 3) × *electrode location* (Fz vs. Cz vs. Pz).

An alpha-level of .05 was defined for all statistical tests. Statistically significant results were reported including the generalized *η*_G_^2^ effect size measure. Greenhouse–Geisser corrections of degrees of freedom were applied where appropriate. Follow up ANOVAs and *t*-tests were computed for statistically significant interactions including two or more factors.

## Results

### Behavioral results

The mean absolute error of reported vs. presented visual targets per block was .6 in the dark condition (SD = .9; min = 0; max = 3.5) and .68 in the light condition (SD = .81; min = 0; max = 3.5). The difference between lighting conditions was not statistically significant (*t*(1,19) = .443; *p* = .663).

**Figure 1.**
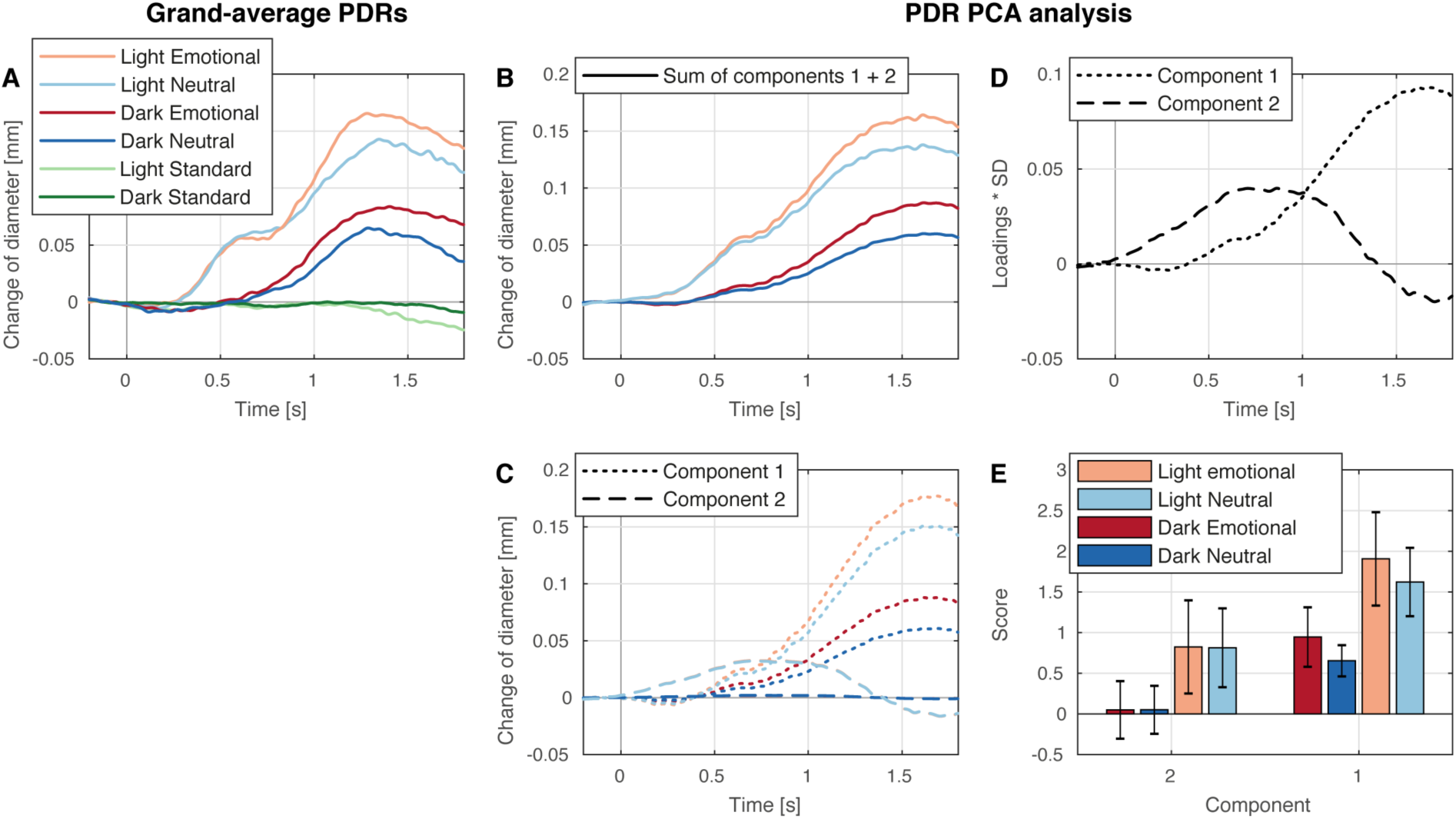
Panel A: Grand-average PDRs. Novels elicited a biphasic PDR. The early peak could be identified in the light condition only. An effect of emotion was only observed for the later peak. Panel D: scaled PCA component time courses (loadings ^*^ SD; mm). Component 2 presumably reflects relaxation of iris sphincter muscle (parasympathetic inhibition). Component 1 presumably reflects constriction of iris dilator muscle (sympathetic activation). Panel E: PCA component scores: Component 2 was only observed in the light condition. An effect of emotion was only observed for component 1. Panel B and C: Reconstructed PDR data per component and summed over both components. The direct comparison of panels A and B shows the fit of the PCA to the observed data (components are ordered by explained variance, component 2 peaks earlier than component 1).

### PDR PCA analysis

A biphasic PDR to novel sounds was observed as reported in previous experiments (Steinhauer & Hakerem, 1992; Wetzel et al., 2016). The early peak could be identified in the light condition only. An effect of emotion was only observed for the later peak. The grand-average PDRs to standard and novel sounds are displayed in Fig. 1A. PCA components loadings and scores are displayed in Fig. 1D–E. Two components were extracted presumably reflecting relaxation of the iris sphincter muscle due to parasympathetic inhibition (component 2; peak latency 0.864 s after sound onset) and constriction of the iris dilator muscle due to sympathetic activation (component 1; peak latency 1.616 s after sound onset) in response to novels. Component scores were significantly higher in the light than in the dark condition (main effect light: *F*(1,19) = 15.262, *p* = .001, *η*_G_^2^ = .192). Note that component 2 could only be observed in the light condition but not in the dark condition (see Fig. 1D). For component 1, scores were significantly higher in response to emotional than in response to neutral sounds but not for component 2 (interaction emotion × component: *F*(1,19) = 5.564, *p* < .026, *η*_G_^2^ = .006; emotional vs. neutral, component 1: *t*(19) = 2.830, *p* = .011; component 2: *t*(19) = .048, *p* = .963). A significant main effects of component (*F*(1,19) = 21.771, *p* < .001, *η*_G_^2^ = .186) was observed but should be interpreted with care as it was included in the interaction effect.

**Figure 2.**
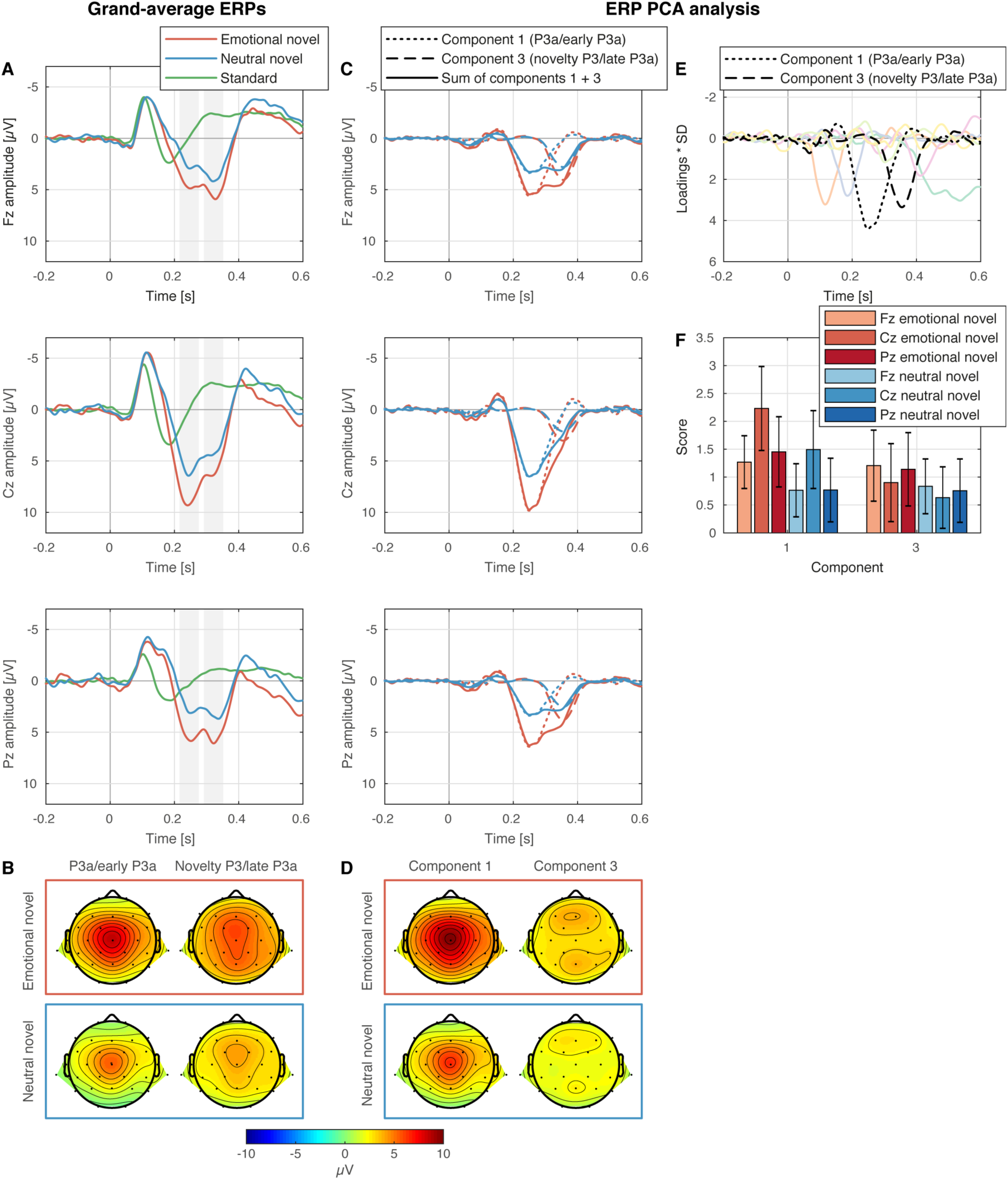
Panel A: Grand-average ERPs. Novels elicited a biphasic P3 response presumably reflecting P3a (early P3a) and novelty P3 (late P3a). Both, P3a and novelty P3 amplitudes were enhanced for emotionally negative sounds. Panel B: Grand-average ERP mean topographies in 60 ms time windows centered around the peaks in the grand-average (242 and 324 ms after sound onset; indicated by grey bars in panel A). Panel E: scaled PCA component time courses (loadings ^*^ SD; µV) for the eight retained components. Panel E: PCA component scores for components 1 (P3a/early P3a) and 3 (novelty P3/late P3a): Both components were enhanced for emotionally negative sounds but had distinct scalp topographies. Panel C and D: Reconstructed ERP data per component and summed over both components and component scalp topographies. Note the considerably distinct scalp topographies of ERP novelty P3 (late P3a) and component 3 indicating substantial contamination of novelty P3 (late P3a) by the overlapping P3a (early P3a) ERP (resolved by PCA).

### ERP PCA analysis

In response to novels sounds an enhanced and delayed N1 response was observed presumably reflecting N1 adaptation and mismatch negativity (MMN) followed by a biphasic P3 component (peaking 242 and 324 ms after sound onset) as reported previously in the literature as P3a and novelty P3 (Barry et al., 2016; Friedman et al., 2001) or early and late P3a (Escera et al., 1998; Yago et al., 2003). Both, P3a and novelty P3 amplitudes were enhanced for emotionally negative sounds. The grand-average ERPs in response to standard and novel sounds are displayed in Fig. 2A–B. PCA components loadings and scores are displayed in Fig. 2E–F. Eight components were extracted. Here, we focus on components 1 (reflecting P3a/early P3a; peak latency 0.248 s after sound onset; central maximum) and 3 (reflecting novelty P3/late P3a; peak latency 0.354 s after sound onset; frontal and parietal maximum). The novelty P3 topography reflects the two spatial frontal and parietal subcomponents previously described by Barry and colleagues (2016). Note that, amplitude, latency, and topography of the novelty P3 (late P3a) component would be substantially biased in a traditional peak centered time window based analysis due the overlap of the two (and other) components. Component scores were significantly modulated by emotion, and electrode location (interaction emotion × component × electrode location: *F*(2,38) = 4.600, *p* < .016, *η*_G_^2^ = .001, ε = .830). For component 1 (P3a/early P3a) the scores were larger for emotional than neutral novels and this difference varied as a function of electrode locations (follow-up interaction emotion × electrode location: *F*(2,38) = 6.711, *p* = .007, *η*_G_^2^ = .002, ε = .781; emotional vs. neutral novel: Fz: *t*(19) = 5.827, *p* < .001; Cz: *t*(19) = 5.874, *p* < .001; *t*(19) = 5.650, *p* < .001). For component 3 (novelty P3/late P3a) the difference in scores between emotional and neutral novels was not significantly different between electrode locations but scores were significantly higher for emotional novels than for neutral novels (follow-up main effect emotion: *F*(1,19) = 5.136, *p* = .035, *η*_G_^2^ = .018). A significant main effect of emotion (*F*(1,19) = 18.296, *p* < .001, *η*_G_^2^ = .036) and significant interactions of emotion and component (*F*(1,19) = 6.098, *p* = .023, *η*_G_^2^ = .004; the effect of emotion was larger for P3a/early P3a, *t*(19) = 6.075, *p* < .001 than for novelty P3/late P3a, *t*(19) = 2.266, *p* < .035), and component and electrode location were observed (*F*(1,19) = 16.184, *p* < .001, *η*_G_^2^ = .035; component scores were significantly different at Cz, *t*(19) = 2.591, *p* < .018, but not at Fz, *t*(19) = 0.011, *p* < .991, and Pz, *t*(19) = 0.484, *p* < .634) but should be interpreted with care as they were also included in the higher-order three-way interaction effect.

## Discussion

In an auditory oddball paradigm task-irrelevant, emotionally arousing negative, and neutral novel sounds were presented amongst frequent sine wave standard sounds in dark and moderate lighting conditions while subjects performed a visual orientation tracking task. A similar pattern of results was observed for the pupil dilation response and the P3 ERP-components: novel sounds elicited a biphasic PDR separable into two components by means of PCA. One component–presumably reflecting parasympathetic inhibition–was absent in darkness, while the other component–presumably reflecting sympathetic activation–was observed in dark and moderate lighting. Only the sympathetic component was enhanced by the emotional content of novel sounds. Novel sounds elicited a biphasic P3 in the ERP. Both, P3a (early P3a) and novelty P3 (late P3a) were enhanced in response to emotional compared to neutral novels.

### Effects of lighting on PDR

As hypothesized by Steinhauer and Hakerem (1992; Marchak & Steinhauer, 2011) the contributions of iris sphincter muscle relaxation and iris dilator muscle constriction to the pupil dilation observed in response to significant, rare, or novel events can indeed be separated by means of PCA. Two components with different time courses could be extracted replicating Wetzel et al. (2016). The fast component was absent in darkness while the slow component was observed in dark and moderate lighting conditions. As the parasympathetic tone is minimal in darkness (Loewenfeld, 1958; and sphincter muscle relaxed) pupil dilation observed in darkness must only be attributed to constriction of the iris dilator muscle due to sympathetic activation. Thus, we could validate the attribution of the faster component (the early part/peak of the biphasic response) to sphincter muscle relaxation due to parasympathetic inhibition and the slower component (the late part/peak of the biphasic response) to dilator constriction due to sympathetic activation previously suggested by Steinhauer and Hakerem (1992). Besides the modulation of the parasympathetic component we also observed a modulation of the sympathetic component by lighting conditions which was not predicted following Steinhauer and colleagues (1992; reflected as main effect of lighting in the ANOVA). We assume that this is due to restricted dynamic range also of the iris dilator muscle in the fully dark-adapted pupil.

### Effects of novels’ emotional arousal on PDR

Novel sounds elicited a PDR presumably signaling surprise and arousal (Einhäuser, 2017; Preuschoff et al., 2011). In line with previous findings we observed larger PDRs to emotionally arousing negative than to neutral novel sounds (Partala & Surakka, 2003). Importantly, a larger PDR to emotionally arousing sounds was only observed for the sympathetic activation component (but not for the parasympathetic inhibition component) of the PDR. The lack of a modulation of the parasympathetic component by emotion is presumably not due to emotional information not being available yet, as the ERP P3a component peaking more than half a second earlier is well affected by sounds induced emotional arousal (see below). Thus, our results give direct evidence for the hypothesis suggested by Bradley et al. (2008), based on indirect evidence, that emotional arousal is specifically reflected in sympathetic nervous system activation. A biphasic PDR separable into two PCA components in response to novel sounds was previously reported for preverbal 14-month-old infants as well as for adults (Wetzel et al., 2016). In their study, the authors reported a significant increase of component scores from the early to the late component in infants for novel baby cry and pink noise sounds. Our present results suggest that this observed modulation of component scores was due to the increased emotional arousal induced by the baby cry and the possibly aversive pink noise sound. Importantly, being able to separate novelty vs. emotional relevance in pre-verbal infants by passive observation of pupil dilation opens promising opportunities in the study of infant cognition

Novel sounds elicited a biphasic P3 component in the ERP. This was frequently described in the literature (Alho et al., 2003; Barry et al., 2016; Escera et al., 1998; Wetzel & Schröger, 2007; Wetzel, Widmann, & Schröger, 2011; Yago et al., 2003). The nomenclature and the functions attributed to the mechanisms underlying each subcomponent are still under debate. Escera et al. (1998) and Alho et al. (2003) labeled them as early and late P3a and suggested that the early peak reflects continued stimulus and deviant feature processing whereas the late peak was suggested to be associated with more general processes involved in the involuntary orienting of attention towards a new unexpected sound (Horvath, Sussman, Winkler, & Schröger, 2011). Yago et al. (2003) attributed both subcomponents to novelty processing and labeled them as early and late novelty P3. In a recently reported series of studies Barry et al. (2016) suggest a distinction of the early peak as genuine P3a indexing attention to the stimulus vs. the late peak as novelty P3 (novelty P3 including two spatial subcomponents) indexing stimulus novelty and the genuine orienting reflex. Here, both P3 component peaks were also separable by means of PCA revealing two sub-components with distinct but overlapping time course and topography. Directly comparing the ERP and PCA P3a (early P3a) and novelty P3 (late P3a) component time course and topographies demonstrates that in particular the novelty P3 ERP peak latency (324 ms ERP peak latency vs. 354 ms PCA component peak latency) and topography (central ERP component peak vs. frontal and parietal PCA component peaks and central trough) is substantially biased by overlap with the larger amplitude P3a (early P3a). Since different cognitive functions are associated with the early and the late subcomponents and assumed underlying sources, this finding should be considered when interpreting modulations of the P3 by experimental conditions in the future

In line with previous results by Thierry and Roberts (2007) the amplitude of both, P3a and novelty P3 (early and late P3a) PCA components was found to be significantly enhanced in response to emotionally arousing compared to neutral novels. The PCA analysis confirms that the enhanced amplitudes are indeed due to a genuine modulation of both components rather than due to an additional emotion sensitive component temporally and spatially overlapping both components. Following the suggested interpretation of P3a and novelty P3 (or early and late P3a) enhanced amplitudes in response to emotionally arousing sounds are supposed to indicate enhanced stimulus-related processing as well as orienting of attention and novel evaluation. It should be noted that Czigler et al. (2007) did not find differences in P3a latency or amplitude for aversive vs. everyday deviant sounds. In this study, sounds were rated for pleasantness (valence) but not for arousal. Thus, it seems possible that the aversive sounds used there did not necessarily induce higher arousal than the everyday sounds

The observed co-modulation of the sympathetic contribution to the PDR and the P3a component is compatible with the hypothesis that the LC-NE system and the peripheral sympathetic nervous system are linked and co-activated by projections from a common medullary pathway (Murphy et al., 2011; for review see Nieuwenhuis et al., 2011). The LC shows a phasic increase of activity in response to salient, unexpected, novel, task-relevant (or otherwise motivationally significant) stimuli leading to NE release in widespread cortical areas increasing cortical neuronal responsivity. In this context, the P3a is considered rather as a neuro-modulatory response allowing the fast evaluation of significant stimuli to prepare for rapid action (Nieuwenhuis et al., 2011; Polich, 2007). Novel sounds are significant as they potentially announce changes in the environment requiring an adaptation of behavior. Emotionally arousing stimuli are additionally associated with intrinsic significance. As a 18 limitation we would like to note that a direct correlation of PDR and novelty P3 amplitudes was not found at single trial level in a previous study by Kamp and Donchin (2015) but only correlations of both P300 and PDR with reaction time and a negative correlation of pupil baseline amplitude (indexing tonic LC activity) with the novelty P3.

In summary, from a functional point of view, surprisingly occurring but task-irrelevant emotionally arousing negative sounds caused enhanced stimulus-related processing involving essential attentional functions. Importantly, we were able to dissociate contributions of parasympathetic inhibition and sympathetic activation to the PDR in response to unexpected task-irrelevant novel sounds. Only the sympathetic contribution and the co-registered P3 components (P3a and novelty P3) were enhanced in response to emotionally arousing compared to neutral novel sounds. This gives direct evidence for the suggestion that emotional arousal is reflected in sympathetic nervous system activation (Bradley et al., 2008) and is compatible with the suggestion that P3 might reflect co-activation of the LC-NE and the sympathetic nervous systems linking the PDR and the P3. The possibility to dissociate parasympathetic and sympathetic activity in the PDR allows promising applications to study processes of attention and cognition in particular in children, patients, or other populations where pupil diameter measurement can be recorded significantly easier than other psychophysiological measures.

## Acknowledgement

We are grateful to N. Coy, F. Haiduk, M. Salditt, E. Schäfer, F. Scharf, T. Scherrer, O. Shevchenko, and B. Sϋßkoch for their help in conducting the experiment. This work was supported by the German Research Foundation (DFG; WE5026/1-1 and WE5026/1-2).

## Reference

Alho, K., Escera, C., & Schröger, E. (2003). Event-related brain potential indices of involuntary attention to auditory stimulus changes. In J. Polich (Ed.), in: Detection of change: Event-related potential and fMRI findings, (pp. 23–40) Dordrecht, Netherlands: Kluwer Academic Publishers.

Aston-Jones, G., & Cohen, J. D. (2005). An integrative theory of locus coeruleus544 norepinephrine function: adaptive gain and optimal performance. Annual Review of Neuroscience, 28, 403–450, doi:10.1146/annurev.neuro.28.061604.135709

Barry, R. J., Steiner, G. Z., & Cohen De Blasio, F. M. (2016). Reinstating the Novelty P3. Scientific Reports, 6, 31200. doi:10.1038/srep31200

Bigdely-Shamlo, N., Mullen, T., Kothe, C., Su, K. M., & Robbins, K. A. (2015). The PREP pipeline: standardized preprocessing for large-scale EEG analysis. Frontiers in Neuroinformatics, 9, 16. doi:10.3389/fninf.2015.00016

Bradley, M.M, Keil, A., & Lang, P. J., (2012). Orienting and emotional perception: facilitation, attenuation, and interference. Frontiers in Psychology, 3, 493 doi:10.3389/fpsyg.2012.00493

Bradley, M.M, & Lang, P. J. (1994). Measuring emotion: the Self-Assessment Manikin and the Semantic Differential. Journal of Behavior Therapy and Experimental Psychiatry, 25, 49–59 doi:10.1016/0005-7916(94)90063-9

Bradley, M.M, & Lang, P. J. (2007). Gainesville, FL: University of Florida. The International Affective Digitized Sounds (2nd ed.; IADS-2): Affective ratings of sounds and instruction manual (Technical Report B-3). Gainesville, FL: University of Florida.

Bradley, M.M, Miccoli, L.Escrig, M. A., & Lang, P. J. (2008). The pupil as a measure of emotional arousal and autonomic activation. Psychophysiology, 45, 602–607 doi:10.1111/j.1469-8986.2008.00654.x

Chaumon, M., Bishop, D. V., & Busch, N. A. (2015). A practical guide to the selection of independent components of the electroencephalogram for artifact correction. Journal of Neuroscience Methods, 250, 47–63 doi:10.1016/j.jneumeth.2015.02.025

Cuthbert, B. N., Schupp, H. T., Bradley, M.M, Birbaumer, N., & Lang, P. J. (2000). Brain potentials in affective picture processing: covariation with autonomic arousal and affective report. Biological Psychology, 52, 95–111

Czigler, I., Cox, T. J., Gyimesi, K., & Horvath, J. (2007). Event-related potential study to aversive auditory stimuli. Neuroscience Lettersw, 420, 251–256 doi:10.1016/j.neulet.2007.05.007

Delorme, A., & Makeig, S. (2004). EEGLAB: an open source toolbox for analysis of single573 trial EEG dynamics including independent component analysis. Journal of Neuroscience Methods, 134, 9–21 doi:10.1016/j.jneumeth.2003.10.009

Delplanque, S., Silvert, L., Hot, P., Rigoulot, S., & Sequeira, H.. (2006). Arousal and valence effects on event-related P3a and P3b during emotional categorization. International Journal of Psychophysiology, 60, 315–322 doi:10.1016/j.ijpsycho.2005.06.006

Delplanque, S., Silvert,L., Hot, P., & Sequeira, H. (2005). Event-related P3a and P3b in response to unpredictable emotional stimuli. Biological Psychology, 68, 107–120. doi:10.1016/j.biopsycho.2004.04.006

Dien, J. (2010). The ERP PCA Toolkit: an open source program for advanced statistical analysis of event-related potential data. Journal of Neuroscience Methods, 187, 138–145. doi:10.1016/j.jneumeth.2009.12.009

Einhäuser, W. (2017). The pupil as marker of cognitive processes. In Q. Zhao (Ed.), Computational and cognitive neuroscience of vision 141&;169;.

Escera, C., Alho, K., Winkler, I., & Näätänen, R. (1998). Neural mechanisms of involuntary attention to acoustic novelty and change. Journal of Cognitive Neuroscience, 10, 590– 604. doi:10.1162/089892998562997

Foti, D., Hajcak, G., & Dien J., (2009). Differentiating neural responses to emotional pictures: evidence from temporal-spatial PCA. Psychophysiology, 46, 521–530.

Friedman, D., Cycowicz, Y. M., & Gaeta H. (2001). The novelty P3: an event-related brain potential (ERP) sign of the brain's evaluation of novelty. Neuroscience and Biobehavioral Reviews, 25, 355–373.

Friedman, D., Hakerem,G., Sutton, S., & Fleiss, J. L. (1973). Effect of stimulus uncertainty on the pupillary dilation response and the vertex evoked potential. Electroencephalography and Clinical Neurophysiology, 34, 475–484. doi:10.1016/0013-4694(73)90065-5

Geuter, S., Gamer, M., Onat, S., & Büchel, C. (2014). Parametric trial-by-trial prediction of pain by easily available physiological measures. Pain, 155, 994–1001. doi:10.1016/j.pain.2014.02.005

Groppe, D. M., Makeig, S., & Kutas, M. (2009). Identifying reliable independent components via split-half comparisons. Neuroimage, 45, 1199–1211. doi:10.1016/j.neuroimage.2008.12.038

Hasting, A., Wassiliwizky, E, & Kotz, S. A. (2010). Erfahrung übertrifft Emotion: Differenzielle Effekte von Vertrautheit und Valenz im ereigniskorrelierten Potential auf Umweltgeräusche im Novelty-Oddball Paradigma. Paper presented at the Psychologie und Gehirn, Greifswald, Germany

Hochmann, J. R., & Papeo, L. (2014). The invariance problem in infancy: a pupillometry study. Psychological Science, 25, 2038–2046. doi:10.1177/0956797614547918

Horvath, J., Sussman, E., Winkler, I., & Schröger, E. (2011). Preventing distraction: assessing stimulus-specific and general effects of the predictive cueing of deviant auditory events. Biological Psychology, 87, 35–48. doi:10.1016/j.biopsycho.2011.01.011

Kamp, S. M., & Donchin, E. (2015). ERP and pupil responses to deviance in an oddball paradigm. Psychophysiology, 52, 460–471. doi:10.1111/psyp.12378

Keil, A., Bradley, M. M., Hauk, O., Rockstroh, B., Elbert, T., & Lang, P. J. (2002). Large617 scale neural correlates of affective picture processing. Psychophysiology, 39, 641–649. doi:10.1017.S0048577202394162

Laeng, B., Sirois, S., & Gredebäck, G. (2012). Pupillometry: A Window to the Preconscious?. Perspectives on Psychological Science, 7, 18–27. doi:10.1177/1745691611427305

Liao, H. I., Kidani, S., Yoneya, M., Kashino, M., & Furukawa, S. (2016). Correspondences among pupillary dilation response, subjective salience of sounds, and loudness. Psychonomic Bulletin and Review, 23, 412–425. doi:10.3758/s13423-015-0898-0

Liao, H. I., Yoneya, M., Kidani, S., Kashino, M., & Furukawa, S. (2016). Human Pupillary Dilation Response to Deviant Auditory Stimuli: Effects of Stimulus Properties and Voluntary Attention. Frontiers in Neuroscience, 10, 43. doi:10.3389/fnins.2016.00043

Loewenfeld, I. E. (1958). Mechanisms of reflex dilatation of the pupil; historical review and experimental analysis. Documenta Ophthalmologica, 12, 185–448. doi:10.1007/BF00913471

Marchak, F. M., & Steinhauer, S. R. (2011). Fundamentals of pupillary measures and eye tracking. Paper presented at the 2011 Annual Meeting of the Society for Psychophysiolocal Research, Boston, MA. http://www.wpic.pitt.edu/research/biometrics/labpubs.html

Max, C., Widmann, A., Kotz, S. A., Schröger, E., & Wetzel, N. (2015). Distraction by emotional sounds: Disentangling arousal benefits and orienting costs. Emotion, 15, 428–437. doi:10.1037/a0039041

McDougal, D. H., & Gamlin, P. D. R. (2008). Pupillary Control Pathways. In A. I. Basbaum, A. Kaneko, G. M. Shepherd, G. Westheimer, T. D. Albright, R. H. Masland, P. Dallos, D. Oertel, S. Firestein, G. K. Beauchamp, M. C. Bushnell, J. H., Kaas, & E. Gardner (Eds.), The Senses: A Comprehensive Reference (Vol. 1, pp. 521–536). San Diego, CA: Academic Press.

Merritt, S. L., Keegan, A. P., & Mercer, P. W. (1994). Artifact management in pupillometry. Nursing Research, 43, 56–59. doi:10.1097/00006199-199401000-00012

Mognon, A., Jovicich, J., Bruzzone, L., & Buiatti, M. (2011). ADJUST: An automatic EEG artifact detector based on the joint use of spatial and temporal features. Psychophysiology, 48, 229–240. doi:10.1111/j.1469-8986.2010.01061.x

Murphy, P. R., O'Connell, R. G., O'Sullivan, M., Robertson, I. H., & Balsters, J. H. (2014). Pupil diameter covaries with BOLD activity in human locus coeruleus. Human Brain Mapping, 35, 4140–4154. doi:10.1002/hbm.22466

Murphy, P. R., Robertson, I. H., Balsters, J. H., & O'Connell, R. G. (2011). Pupillometry and P3 index the locus coeruleus-noradrenergic arousal function in humans. Psychophysiology, 48, 1532–1543. doi:10.1111/j.1469-8986.2011.01226.x

Nieuwenhuis, S., De Geus, E. J., & Aston-Jones, G. (2011). The anatomical and functional relationship between the P3 and autonomic components of the orienting response. Psychophysiology, 48, 162–175. doi:10.1111/j.1469-8986.2010.01057.x

Pakarinen, S., Sokka, L., Leinikka, M., Henelius, A., Korpela, J., & Huotilainen, M. (2014). Fast determination of MMN and P3a responses to linguistically and emotionally relevant changes in pseudoword stimuli. Neuroscience Letters, 577, 28–33. doi:10.1016/j.neulet.2014.06.004

Partala, T., & Surakka, V. (2003). Pupil size variation as an indication of affective processing. International Journal of Human-Computer Studies, 59, 185–198. doi:10.1016/S1071-5819(03)00017-X

Polich, J. (2007). Updating P300: an integrative theory of P3a and P3b. Clinical Neurophysiology, 118, 2128–2148. doi:10.1016/j.clinph.2007.04.019

Preuschoff, K., ’t Hart, B. M., & Einhäuser, W. (2011). Pupil Dilation Signals Surprise: Evidence for Noradrenaline's Role in Decision Making. Frontiers in Neuroscience, 5, 115. doi:10.3389/fnins.2011.00115

Qiyuan, J., Richer, F., Wagoner, B. L., & Beatty, J. (1985). The pupil and stimulus probability. Psychophysiology, 22, 530–534. doi:10.1111/j.1469-8986.1985.tb01645.x

Sara, S. J., & Bouret, S. (2012). Orienting and reorienting: the locus coeruleus mediates cognition through arousal. Neuron, 76, 130–141. doi:10.1016/j.neuron.2012.09.011

Steinhauer, S. R., & Hakerem, G. (1992). The pupillary response in cognitive psychophysiology and schizophrenia. Annals of the New York Academy of Sciences, 658, 182–204. doi:10.1111/j.1749-6632.1992.tb22845.x

Steinhauer, S. R., Siegle, G. J., Condray, R., & Pless, M. (2004). Sympathetic and parasympathetic innervation of pupillary dilation during sustained processing. International Journal of Psychophysiology, 52, 77–86. doi:10.1016/j.ijpsycho.2003.12.005

Thierry, G., & Roberts, M. V. (2007). Event-related potential study of attention capture by affective sounds. Neuroreport, 18, 245–248. doi:10.1097/WNR.0b013e328011dc95

Wetzel, N. (2015). Effects of the short-term learned significance of task-irrelevant sounds on involuntary attention in children and adults. International Journal of Psychophysiology, 98, 17–26. doi:10.1016/j.ijpsycho.2015.06.003

Wetzel, N., Buttelmann, D., Schieler, A., & Widmann, A. (2016). Infant and adult pupil dilation in response to unexpected sounds. Developmental Psychobiology, 58, 382– 392. doi:10.1002/dev.21377

Wetzel, N., & Schröger, E. (2007). Modulation of involuntary attention by the duration of novel and pitch deviant sounds in children and adolescents. Biological Psychology, 75, 24–31. doi:10.1016/j.biopsycho.2006.10.006

Wetzel, N., Widmann, A., & Schröger, E. (2011). Processing of novel identifiability and duration in children and adults. Biological Psychology, 86, 39–49. doi:10.1016/j.biopsycho.2010.10.005

Widmann, A., & Schröger, E. (2012). Filter effects and filter artifacts in the analysis of electrophysiological data. Frontiers in Psychology, 3, 233. doi:10.3389/fpsyg.2012.00233

Widmann, A., Schröger, E., & Maess, B. (2015). Digital filter design for electrophysiological data––a practical approach. Journal of Neuroscience Methods, 250, 34–46. doi:10.1016/j.jneumeth.2014.08.002

Winkler, I., Debener, S., Müller, K. R., & Tangermann, M. (2015). On the influence of high700 pass filtering on ICA-based artifact reduction in EEG-ERP. Conference Proceedings of the Annual International Conference of the IEEE Engineering in Medicine and Biology Society, 2015, 4101–4105. doi:10.1109/EMBC.2015.7319296

Yago, E., Escera, C., Alho, K., Giard, M. H., & Serra-Grabulosa, J. M. (2003). Spatiotemporal dynamics of the auditory novelty-P3 event-related brain potential. Brain Research: Cognitive Brain Research, 16, 383–390. doi:10.1016/S0926-6410(03)00052-1

